# Systemic administration of Ivabradine, an HCN channel inhibitor, blocks spontaneous absence seizures

**DOI:** 10.1101/2021.02.17.431588

**Authors:** Yasmine Iacone, Tatiana P. Morais, François David, Francis Delicata, Joanna Sandle, Timea Raffai, H. Rheinallt Parri, Johan Juhl Weisser, Christoffer Bundgaard, Ib Vestergaard Klewe, Gábor Tamás, Morten Skøtt Thomsen, Vincenzo Crunelli, Magor L. Lőrincz

## Abstract

**Objective:** Hyperpolarization-activated cyclic nucleotide-gated (HCN) channels are known to be involved in the generation of absence seizures (ASs), and there is evidence that cortical and thalamic HCN channel dysfunctions may have a pro-absence role. Many HCN channel blockers are available, but their role in ASs has been investigated only by localized brain injection or in *in vitro* model systems due to their limited brain availability. Here, we investigated the effect on ASs of orally administered ivabradine (an HCN channel blocker approved for the treatment of heart failure in humans) following injection of the P-glycoprotein inhibitor elacridar, that is known to increase penetration into the brain of drug substrates for this efflux transporter. The action of ivabradine was also tested following *in vivo* microinjection in the cortical initiation network (CIN) of the somatosensory cortex and in the thalamic ventrobasal nucleus (VB) as well as on cortical and thalamocortical neurons in brain slices.

**Methods:** We used EEG recordings in freely moving Genetic Absence Epilepsy from Strasbourg Rats (GAERS) to assess the action of oral administration of ivabradine, with and without elacridar, on ASs. Ivabradine was also microinjected in the CIN and VB of GAERS *in vivo* and applied to Wistar CIN and GAERS VB slices while recording patch-clamped cortical layer 5/6 and thalamocortical neurons, respectively.

**Results:** Oral administration of ivabradine markedly and dose-dependently reduced ASs. Ivabradine injection in CIN abolished ASs and elicited small-amplitude 4-7 Hz waves (without spikes), whereas in the VB it was less potent. Moreover, ivabradine applied to GAERS VB and Wistar CIN slices selectively decreased HCN-channel-dependent properties of cortical layer 5/6 pyramidal and thalamocortical neurons, respectively.

**Significance:** These results provide the first demonstration of the anti-absence action of a systemically administered HCN channel blocker, indicating the potential of this class of drugs as a novel therapeutic avenue for ASs.

## Introduction

Absence seizures (ASs) are characterized by loss of consciousness and lack of voluntary movements accompanied by generalized spike-and-wave discharges (SWDs) in the electroencephalogram (EEG). ASs are present in several epilepsies and are the only clinical symptom of Childhood Absence Epilepsy (CAE),^1,2^ which accounts for 10-17% of all children with epilepsy^3^ and carries a burdensome personal, familial and societal impact.^4,5^ The first-line therapy for CAE is ethosuximide monotherapy, followed by valproic acid or lamotrigine,^6^ but about 30% of CAE children are pharmaco-resistant, resulting in polytherapy and a consequent marked increase in adverse effects.^7–9^ Furthermore, about 60% of children with AS experience neuropsychiatric comorbidities (mainly attention/cognitive impairments), which can be present before the epilepsy diagnosis, and even be aggravated following full pharmacological control of ASs.^10,11^ Hence, there is a pressing need to find new effective targets for AS treatment.

The role for hyperpolarization-activated cyclic nucleotide-gated (HCN) channels in ASs has been extensively investigated, with many studies reporting a gain- or loss-of-function, mainly involving HCN1 and HCN2 subtypes. In particular, analysis of recombinant HCN2 variants in humans with febrile seizures and genetic epilepsy with febrile seizures-plus shows an increased hyperpolarization-activated current (I_h_), the current generated by HCN channels.^12^ Moreover, several HCN1 variants have been identified in children with early infantile epileptic encephalopathy that lead both to a gain- or loss-of-function,^13^ while in sporadic idiopathic generalized epilepsy patients, point mutations of HCN2 give rise to a channel loss-of-function.^14^ The diversity of these HCN channel dysregulations, however, together with the fact that ASs are not the only phenotype present in these patient cohorts, makes it difficult to draw any firm conclusion on the precise role of HCN channels in human ASs.

Studies in experimental animals have also demonstrated a critical role for HCN channels in ASs, but the results are often contradictory. Global knock-out of HCN1^15^ or HCN2^16^ elicits ASs, suggesting an anti-absence role of these channels. Furthermore, two genetic AS models showed both an increased thalamic and a decreased cortical I_h_ as well as a contrasting up- or down-regulation of HCN1 in both thalamus and cortex,^17–20^ two brain regions that are essential for AS generation.^21,22^ Moreover, removal of the cAMP-sensitivity of HCN2 in the whole brain or thalamic ventrobasal nucleus (VB)-selective HCN2 knock-down lead to ASs,^23^ while VB-selective HCN4 knock-down has no effect.^24^ Conversely, pharmacological and genetic suppression of HCN channels in the VB suppress ASs in three animal models, suggesting that HCN channels in this brain region have a pro-absence role.^25^

Notwithstanding the therapeutic potential of targeting HCN channel for the treatment of ASs, an HCN channel modulator must show efficacy following systemic administration for it to have clinical applicability. Several HCN channel blockers have been developed so far,^26^ including ivabradine (IVA) (Procoralan^©^, Corlanor^©^), a drug approved for the treatment of heart failure.^27–29^ All these HCN channel blockers, however, show a very limited ability to cross the blood-brain barrier (BBB), due to the efflux mediated by P-glycoproteins (Pgp). Thus, whereas HCN channel blockers have been extensively used to inhibit neuronal HCN channels *in vitro*,^30^ the interpretation of the few brain investigations that used systemic administration are questionable due to their poor brain penetration.^31–33^

Here, we show for the first time that IVA orally administered together with Elacridar (ELA), a Pgp inhibitor,^34^ elicits a marked and long-lasting reduction of ASs in a well-established AS model (the Genetic Absence Epilepsy Rats from Strasbourg, GAERS).^35^ Additionally, a similar anti-absence action occurs when IVA is directly injected in the cortical initiation network (CIN)^36^ and the VB, and IVA selectively decreases HCN channel-dependent properties of cortical layer 5/6 pyramidal and VB thalamocortical neurons *in vitro*.

## Materials and Methods

### Animals

GAERS (originally obtained from Strasbourg, France) were bred and maintained at Cardiff University (UK) and University of Szeged (Hungary). Wistar rats (originally from Envigo) were maintained at H. Lundbeck A/S (Valby, Denmark) and University of Szeged (Szeged, Hungary). Animals were provided with normal diet and water *ad libitum*, and kept under a light:dark cycle of 12:12 hours with light on at 7:00 AM. Experimental procedures were performed in agreement with the UK Animals (Scientific Procedures) Act (1986), the European Communities Council Directive (2010/63/EU) and the Danish legislation (Law and Order on Animal experiments; Act No. 474 of 15/05/2014 and Order No. 12 of 07/01/2016).

### IVA plasma and brain bioanalysis

Plasma and brain concentrations of IVA were measured in 3 month-old male Wistar rats 1h after injection to assess its concentration at the time of the ELA peak brain concentration, and in 3-4 month old male GAERS 2h after the injection (i.e. at the end of the recording session) to measure brain IVA concentration at time of the last recording period (see below). Blood samples were kept in 1.6 mg EDTA/ml of blood, centrifuged at 3300 g for 15 min at 4°C and stored at −80°C until bioanalysis. Brains were stored at −80°C until bioanalysis.

Brain samples were prepared by homogenizing half brain using isothermal focused acoustic ultrasonication (Covaris E220x, Covaris, Inc., Woburn, MA) as described previously.^37^ After homogenization calibration standards and QC samples were prepared in blank rat plasma and brain homogenate in a range of 0.5 ng/ml to 500 ng/ml. On the day of analysis, 25 µl of brain homogenates, plasma samples, calibration standards and QC samples were precipitated with 100 µl acetonitrile containing internal standard. After centrifugation (20 min at 3500 g) 50 µl supernatant was transferred to a 300 µl 96-well plate and mixed with 150 µl 25 % acetonitrile solution.

Ivabradine concentrations were determined using ultraperformance liquid chromatography (Acquity UPLC system; Waters, Milford, MA) coupled to a tandem mass spectrometry detector (Waters Xevo TQXS). Chromatographic separation was performed on a Waters C18SB HSS column (30 x 2.1 mm, 1.8 μm particles) with a column compartment temperature of 40°C using gradient elution with mobile phases consisting of 0.1 % formic acid in water and 0.1 % formic acid in acetonitrile. Autosampler temperature was 10°C and injection volume was 5 µl. Electrospray ionization was performed in positive mode. For Ivabradine the ion 469.3^+^ → 262.1^+^ was monitored.

### Surgical procedures

Adult male GAERS (250-300 g) were anaesthetized with isoflurane (2-5%) and body temperature maintained at 37°C with a heating pad. Six gold-plated epidural screws (Svenska Dentorama AB, Sweden) were implanted in pairs, in frontal, parietal and cerebellar sites. For microinjection in the VB and CIN, one 8 mm long guide-cannula (C315G/50-99, Bilaney, UK) was also implanted in both hemispheres (VB: AP-3.2, ML+/- 3.6, DV4.5, with a 5° angle; CIN: AP-2.52, ML+/-4.8, DV1.3.^38^ The animals were allowed to recover for at least five days prior to experiments.

### EEG recordings

The day before experiments, rats were connected to the recording apparatus and placed individually in a plexiglass box within a Faraday cage for 1-2 hours habituation. On the day of the recordings, animals were placed into the plexiglass box for 30 minutes (habituation period) followed by 40 minutes recording (control period). They were then transferred to an anaesthesia induction chamber, slightly anaesthetized with isoflurane (1%) and injected intravenously with either ELA (5mg/kg, 5ml/kg) or vehicle (VEH1) (hydroxypropyl-β-cyclodextrin) as soon as the righting reflex was lost. Anaesthesia was then terminated, and 20 min after the intravenous ELA (or VEH1) administration, the animals received either IVA (10, 20 and 30 mg/kg, all 10 ml/kg) or vehicle (VEH2) (5% D-glucose in distilled water) orally. Rats were then placed in the plexiglass box, reconnected to the EEG apparatus and recorded for 2 hours while being continuously monitored by one researcher. Drug-treatments (VEH1-VEH2, ELA-VEH2, VEH1-IVA and ELA-IVA) were assigned in a pseudo-random manner with a cross-over design. Each animal received a maximum of four different treatments with at least five days between each treatment.

For microinjection experiments, on the day of the experiment the animals were recorded for 1h (control period), followed by the bilateral insertion of a 9 mm (C315I/20-49, Bilaney) cannula, that was connected to a micropump (CMA 400, Linton Instruments, UK). Animals were then injected with either artificial cerebrospinal fluid (aCSF) or IVA (6 nmoles) using a flow rate of 0.25 µl/min for 2 min and recorded for 2 hours.

To check cannulae position, brains were collected and washed in PBS (10 mM) followed by fixation in 4% PFA for 24h. After fixation, 100 µm coronal slices were cut from the region containing the CIN or VB and then mounted with VECTASHIELD® Antifade Mounting Media (Vector Laboratories, UK). Slices were imaged within the next 24 hours and photographed with an Olympus BX61 microscope (Olympus, Japan) with a 4X objective. Results from animals with misplaced cannulae position were not included in the final analysis.

### Data acquisition and analysis SWD detection

The analog EEG signal was acquired through a 4-channels differential pre-amplifier (high-pass filter 0.1 Hz, SuperTech, Hungary) connected to a 4-channel BioAmp amplifier (1000 gain, low-pass filter 500 Hz, SuperTech, Hungary) and digitized at 1000 Hz using a CED Mk3 1401 (Cambridge Electronic Design, UK). SWDs were initially detected semi-automatically using the SeizureDetect script (kindly provided by Steve Clifford, CED) in Spike2 v7.03 (Cambridge Electronic Design, UK), and then checked by visual inspection. Data were digitally processed and an interictal EEG period of wakefulness was manually selected and used to set a threshold of +/- 5-8 SD of the baseline EEG. To identify SWDs, all crossings above or below the threshold were then grouped into bursts according to five pre-set parameters: a maximum onset interval (0.2s), a maximum interval (0.75 s), a minimum number of spikes (5), a minimum interval within bursts (1 s) and a minimum duration (0.6 s). Identified bursts lasting less than 1 second were discarded. The putative bursts were ultimately classified into SWDs according to the frequency, which was manually set between 5 and 12 Hz to exclude deep sleep epochs or artefacts. This semi-automatic detection was further refined by visual inspection. The following parameters were extracted from the EEG data in 20 min epochs: the total time spent in seizure, the total number of seizures and the average duration of a seizure. Treatment data were normalized to the respective control period and statistical analysis performed after normalization (see Statistical Analysis).

### Power spectral analysis

The Welch power spectral density analysis was performed with MATLAB (R2019a, MathWorks, USA) on interictal EEG periods which were devoid of ASs in GAERS treated oral administration of IVA and other drug combinations. Change of power spectrum density between control and treatment periods were measured as baseline percentage. Statistical analysis was performed after normalization to the respective control period (see Statistical Analysis). Similar power spectra were performed on the EEG of GAERS that received IVA directly in the CIN and VB, and compared to VEH injection.

### Cortical and thalamic slice preparation, whole-cell recordings and data analysis

Male Wistar and GAERS rats (both 25-35 days-old) were anesthetized (ketamine/xylazine: 80/8 mg/kg), their brains quickly sliced (320 µm thickness) in the coronal plane, and slices containing either the CIN or the VB were incubated at room temperature (20°C) in aCSF containing (in mM): 130 NaCl, 3.5 KCl, 1 NaH_2_PO_4_, 24 NaHCO_3_, 1 CaCl_2_, 3 MgSO_4_, 10 glucose. For recording, slices were submerged in a chamber perfused with a warmed (35°C) continuously oxygenated (95% O_2_, 5% CO_2_) aCSF containing (in mM): 130 NaCl, 3.5 KCl, 1 KH_2_PO_4_, 24 NaHCO_3_, 1 MgSO_4_, 2 CaCl_2_, and 10 glucose.

Whole-cell patch-clamp recordings were performed using a Heka EPC9 amplifier (Heka Elektronik). Patch pipettes (tip resistance: 4–5 MΩ) were filled with an internal solution containing the following (in mM): 126 K-gluconate, 4 KCl, 4 ATP-Mg, 0.3 GTP-Na_2_, 10 HEPES, 10 kreatin-phosphate (pH 7.25, osmolarity 275 mOsm. The liquid junction potential (−13 mV) was corrected offline. Access and series resistances were constantly monitored, and data from neurons with a >20% change from the initial value were discarded. Action potential amplitude was measured from threshold (20 mV/ms on the first derivative of the membrane potential) to the peak of the action potential. Analysis of these whole-cell data was performed using custom routines written in Igor.

### Statistical analysis

Statistical analysis was performed using GraphPad Prism version 9.0 (GraphPad Software, San Diego, USA). Normality of the data was verified using the QQ plot. The comparison between the two doses of ELA was performed using an unpaired t-test assuming equal variances between the two groups. The effect of each treatment on SWDs following systemic injection was assessed by repeated-measurements (RM) two-way-Analysis of Variance (ANOVA) using Sidak correction for multiple comparisons. The main effect of treatment vs vehicle was also measured as area-under-the-curve (AUC) and analysed with one-way-ANOVA using Dunnett’s multiple comparisons test. Statistical analysis of the power spectra was carried out with one-sided Wilcoxon test comparing treatment with control. The main effect of IVA vs aCSF for CIN and VB microinjections was assessed by two-way ANOVA for multiple timepoints and unpaired *t*-test for AUC. Data from *in vitro* recordings in cortical and thalamic neurons were analysed with Wilcoxon signed ranks test.

All quantitative data in the text and figures are reported as mean+/-SEM, unless stated otherwise.

### Drugs

IVA (3-[3-({[(7S)-3,4-dimethoxybicyclo[4.2.0]octa-1,3,5-trien-7-yl]methyl}(methyl) amino) propyl]-7,8-dimethoxy-2,3,4,5-tetrahydro-1H-3-benzazepin-2-one hydrochloride)) and ELA (N-[4-[2-(3,4-Dihydro-6,7-dimethoxy-2(1H)-isoquinolinyl)ethyl]phenyl]-9,10-dihydro-5-methoxy-9-oxo-4-acridinecarboxamide) were purchased from Sigma-Aldrich. For systemic injections, IVA was dissolved in a 5% D-glucose (Sigma Aldrich) solution and the pH adjusted to 4. ELA was dissolved in 10% Hydroxypropyl-β-cyclodextrin and the pH adjusted to 4. For local microinjections, IVA was diluted in aCSF. All drugs were freshly prepared on each day of experiments.

## Results

### Systemic injection of IVA

The highest dose of IVA (30 mg/kg) used in this study was selected on the basis of its efficacy and safety as described in previous EMA and FDA reports.^27,28^ For selecting a dose of ELA that could lead to suitable brain levels of IVA, we tested two doses of ELA that had been previously reported to allow good brain penetration of other systemically dosed Pgp substrates.^39^ Intravenous pre-treatment of Wistar rats with 5 mg/kg ELA provided higher brain concentration of orally administered IVA (622 ±107 ng/g,) compared to 2.5 mg/kg ELA (259 ± 57.6 ng/g, p=0.018) (**Figure 1A**). Similarly, the brain free concentration of IVA was higher in animals pre-treated with 5 mg/kg than 2.5 mg/kg ELA (307 ± 53.0 and 128 ± 28.4 nM, respectively, p=0.018), showing a direct correlation with the total drug concentration (**Figure 1B**). No significant differences in the IVA plasma levels were observed in animals pre-treated with 2.5 and 5 mg/kg ELA (p=0.603) (**Figure 1A, B**). A dose of 5 mg/kg of ELA was thus used in further experiments.

**Figure 1.**
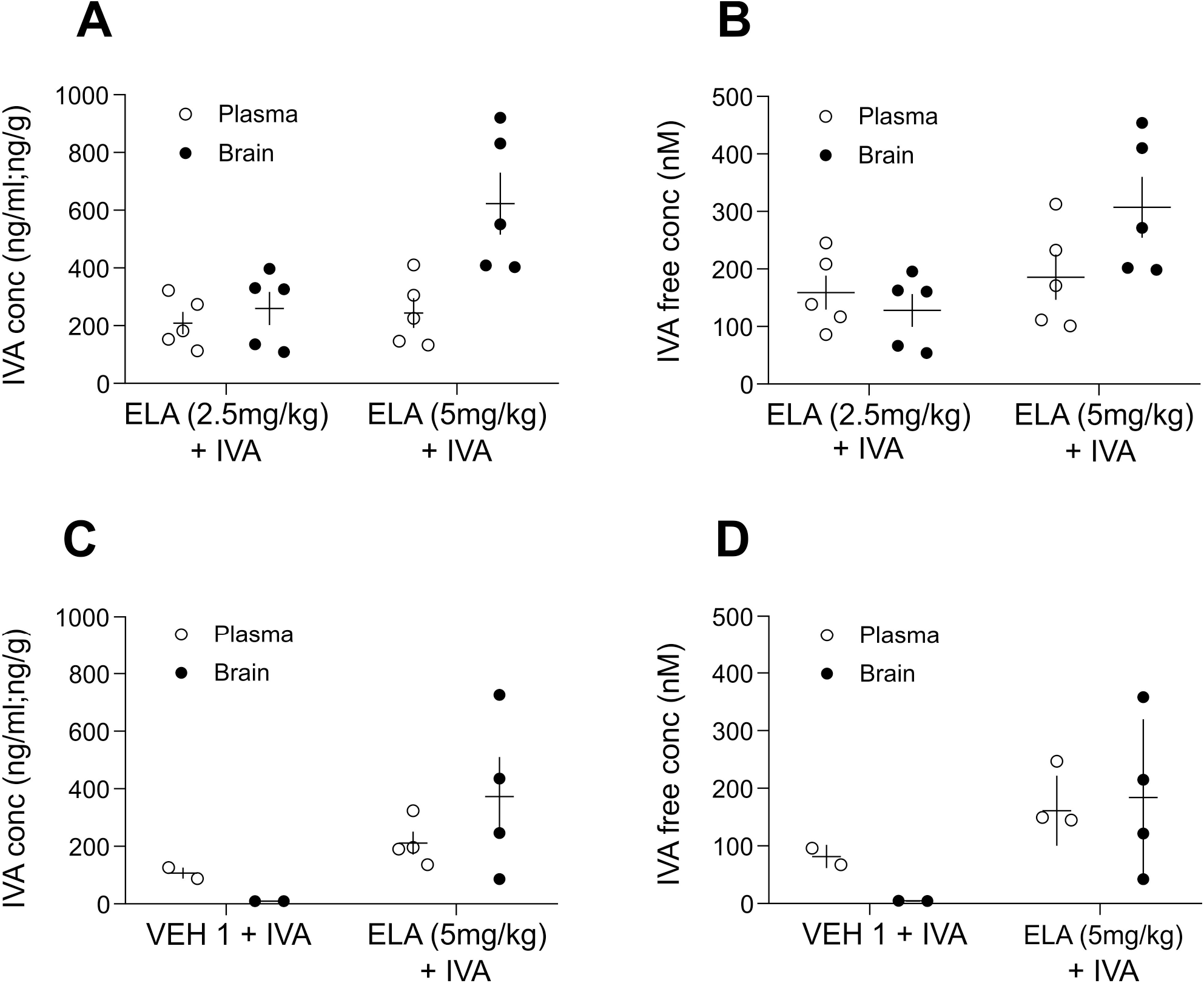
Brain and plasma levels of systemically injected ivabradine with and without pre-treatment with elacridar. **A,B**. Total brain (black dots) and plasma (white dots) levels (**A**) and brain and plasma free concentrations (**B**) measured 1 hour after oral administration of Ivabradine (IVA) (30 mg/kg) in Wistar rats (N=5 each group) that had been pre-treated with intravenous injection of either 2.5 or 5 mg/kg Elacridar (ELA) 30 min before IVA injection. **C,D**. Total brain (black dots) and plasma (white dots) levels (**C**) and brain and plasma free concentrations (**D**) of IVA in GAERS rats treated with VEH1+IVA (N=2) or ELA (5mg/kg) + IVA (N=4). Animals were sacrificed at the end of the EEG recordings, i.e. 2 hours after IVA injection. In **A**-**D**, horizontal and vertical lines indicate mean and ±SEM, respectively.

We next investigated the effect of orally administered IVA on spontaneous ASs in freely moving GAERS. No gross behavioural changes were observed in any treatment group during the EEG recordings described below. As shown in **Figure 2A**, in animals pre-treated with 5 mg/kg ELA, oral administration of 20 and 30 (but not 10) mg/kg IVA markedly reduced spontaneous ASs, an effect that for the highest dose was visible as early as 20 min after IVA administration and lasted for the entire duration of the recorded period (2 hours). Statistical analysis of the normalized area-under-the-curve (AUC) of the entire test period showed a significant reduction (62%) of the total time spent in seizures for the ELA+IVA30 group (0.38±0.09, F=8.77, DFn=5, DFd=22, p=0.0044), but not for the ELA+IVA20 (0.53±0.09,p=0.07), ELA+IVA10 (1.05±0.09, p>0.99), ELA+VEH2 (1.2±0.16, p=0.61) and VEH1+IVA groups (1.12±0.11, p=0.86) compared to VEH1+VEH2 (**Figure 2B1**). The mean duration of the seizures was also significantly decreased (F=6.67, DFn=5, DFd=22, 41%) in the ELA+IVA30 group (0.59±0.07, p=0.008), but not for ELA+IVA20 (0.66±0.09, p=0.09), ELA+IVA10 (1.03±0.1, p>0.99), ELA+VEH2 (1.06±0.08, p>0.99) and VEH1+IVA groups (1.06±0.09, p=0.95) compared to VEH1+VEH2 (**Figure 2B2**). Moreover, the number of seizures showed a significant decrease (45%) in rats treated with ELA+IVA30 (0.55±0.09, F=4.35, DFn=5, DFd=46, p=0.01) but not for ELA+IVA20 (0.71±0.09, p=0.24), ELA+IVA10 (0.99±0.09, p>0.99), ELA+VEH2 (1.08±0.13, p=0.98) and VEH1+IVA groups (1.03±0.06, p=0.99) compared to VEH1+VEH2 (**Figure 2B3**). Finally, power spectra of the interictal EEG showed the administration of ELA+IVA30 to elicit a significant increase in the power of theta (4-8 Hz) and low gamma (30-50 Hz) frequency bands compared to the VEH1+VEH2 group and a small decrease in the alpha band (8-14 Hz) (**Figure 2C**).

**Figure 2.**
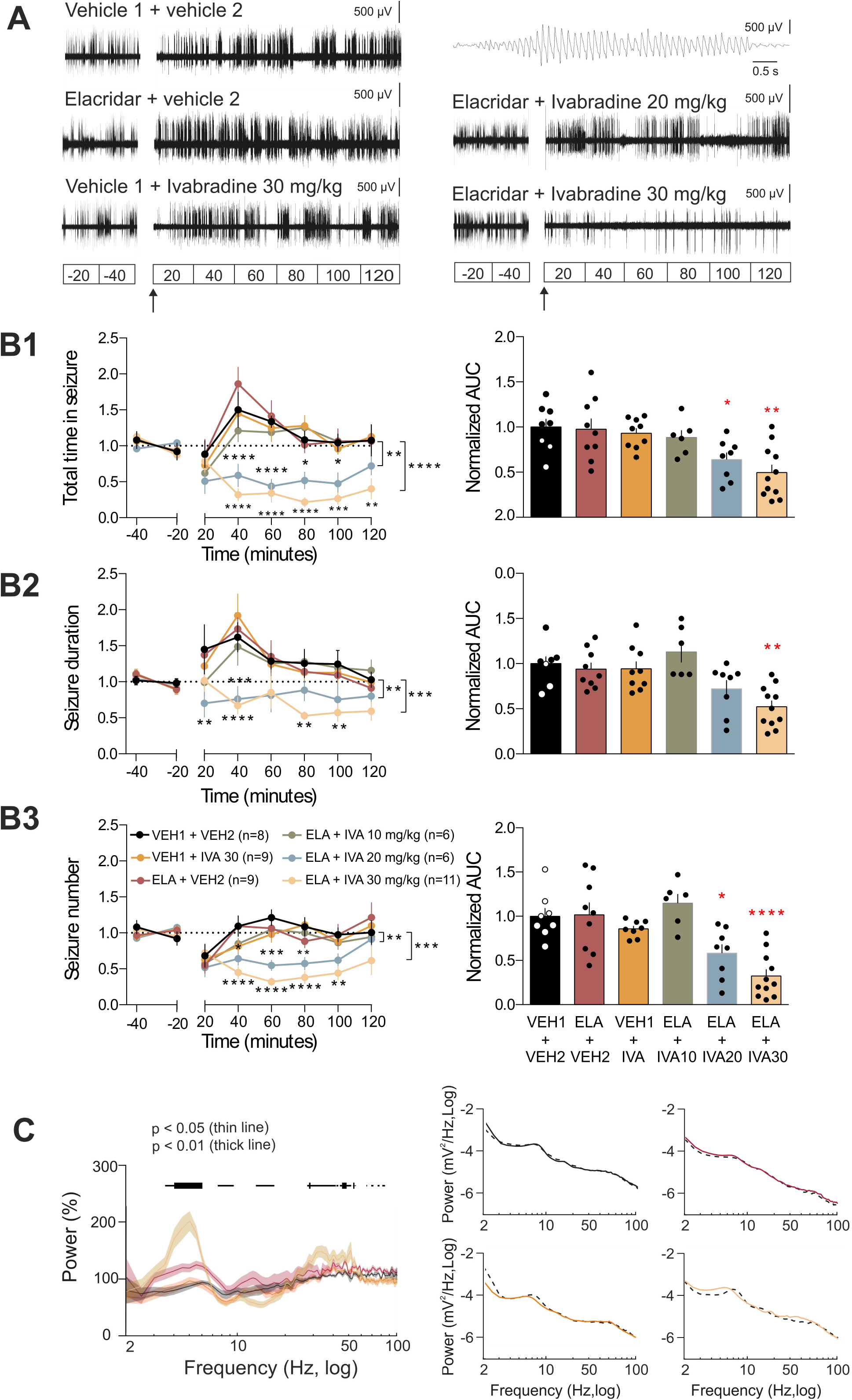
Oral administration of ivabradine markedly blocks ASs. **A**. Representative EEG traces of four different freely moving GAERS injected with either VEH1+VEH2, VEH1+IVA, ELA+VEH2, ELA+IVA10, ELA+IVA20 and ELA+IVA30. Note the marked reduction in SWDs in the ELA+IVA30 group compared to all other groups (a typical SWD is shown enlarged in the top trace on the right). In **A** and the left plots of **B1-3**, interruption in the traces is due to the animals being disconnected from the EEG wires for drug administration: ELA (or VEH1) was injected at time −20 min and IVA (or VEH2) at time 0 (indicated by the black arrow). Data in **B1-3** are normalized to the control period (see methods). **B1**. Time-response curves (left graph) and area under the curve (AUC) of the whole treatment period (right graph) of the total time spent in seizures for VEH1+VEH2 (black, N=9), ELA+VEH2 (red, N=9), VEH1+IVA (orange, N=9), ELA+IVA10 (green, N=6), ELA+IVA20 (blue, N=8) and ELA+IVA (ochre, N=11) groups (*p<0.05, **p < 0.01, ***p<0.005, ****p<0.001) (colour-code and number of animals as in **B3**). **B2**. Time-response curve (left graph) and AUC (right graph) of the whole treatment period of seizure duration for the six treatment groups (*p<0.05, **p < 0.01, ***p<0.005, ****p<0.001). **B3**. Time-response curve (left graph) and AUC (right graph) of the whole treatment period of the number of seizures for the six treatment groups (*p<0.05, **p < 0.01, ***p<0.005, ****p<0.001). **E**. Left graph: Average normalized interictal power spectra of the four treatment groups (number of animals as in **B3**, except VEH1+VEH2 N=6). Horizontal bars on top indicate statistical significance (thin line: p < 0.05; thick line: p < 0.01). Right: representative examples of interictal power spectra for individual animals of the VEH1+VEH2 (black), ELA+VEH2 (red), VEH1+IVA (orange) and ELA+IVA30 (ochre) groups showing both the control period (dashed black line) and the treatment period.

At the end of the last recording session, the brain and plasma of those rats that had received ELA+IVA30 or VEH+IVA30 as their last treatment were collected to determine the IVA plasma and brain levels. As shown in **Figure 1C**, the total plasma concentrations of IVA measured 2 hours after dosing were in the same range in the two treatment groups (VEH1+IVA30: 107 ± 19 ng/ml; ELA+IVA30 212 ± 39 ng/ml) while the brain concentration was substantially higher in the animals that were dosed with ELA-IVA30 (373 ± 137 ng/g) compared to those with VEH1+IVA30 (9.5 ± 0.3 ng/g). The free brain concentration of IVA was 4.7 ± 0.1 nM for VEH1+IVA30 and 184 ± 67.8 nM for the ELA+IVA30 group (**Figure 1D**).

### Local microinjection of IVA

Since ASs are generated by abnormal firing in cortico-thalamo-cortical circuits, we next investigated whether the anti-absence effect of systemically administered IVA was mediated by an action on thalamic and/or cortical regions. Thus, we applied IVA by bilateral microinjection in the VB or CIN of freely moving GAERS.

Bilateral microinjection of IVA (6 nmoles) in the VB of freely moving GAERS reduced ASs compared to VEH injection (**Figure 3A**). Statistical analysis of the AUC of the entire test period showed a significant reduction (40%, F=2.61, DFn=6, DFd=4, p=0.05) of the total time spent in seizures of IVA (0.6±0.09) compared to VEH (**Figure 3B1**). The number of seizures decreased (29%, F=2.99, DFn=6, DFd=4, p=0.011) in rats treated with IVA (0.71±0.05) compared to VEH (**Figure 3B2**), but the mean duration of seizures was unchanged (5%, F=1.39, DFn=6, DFd=4, p=0.72) (0.95±0.12) (**Figure 3B3**).

**Figure 3.**
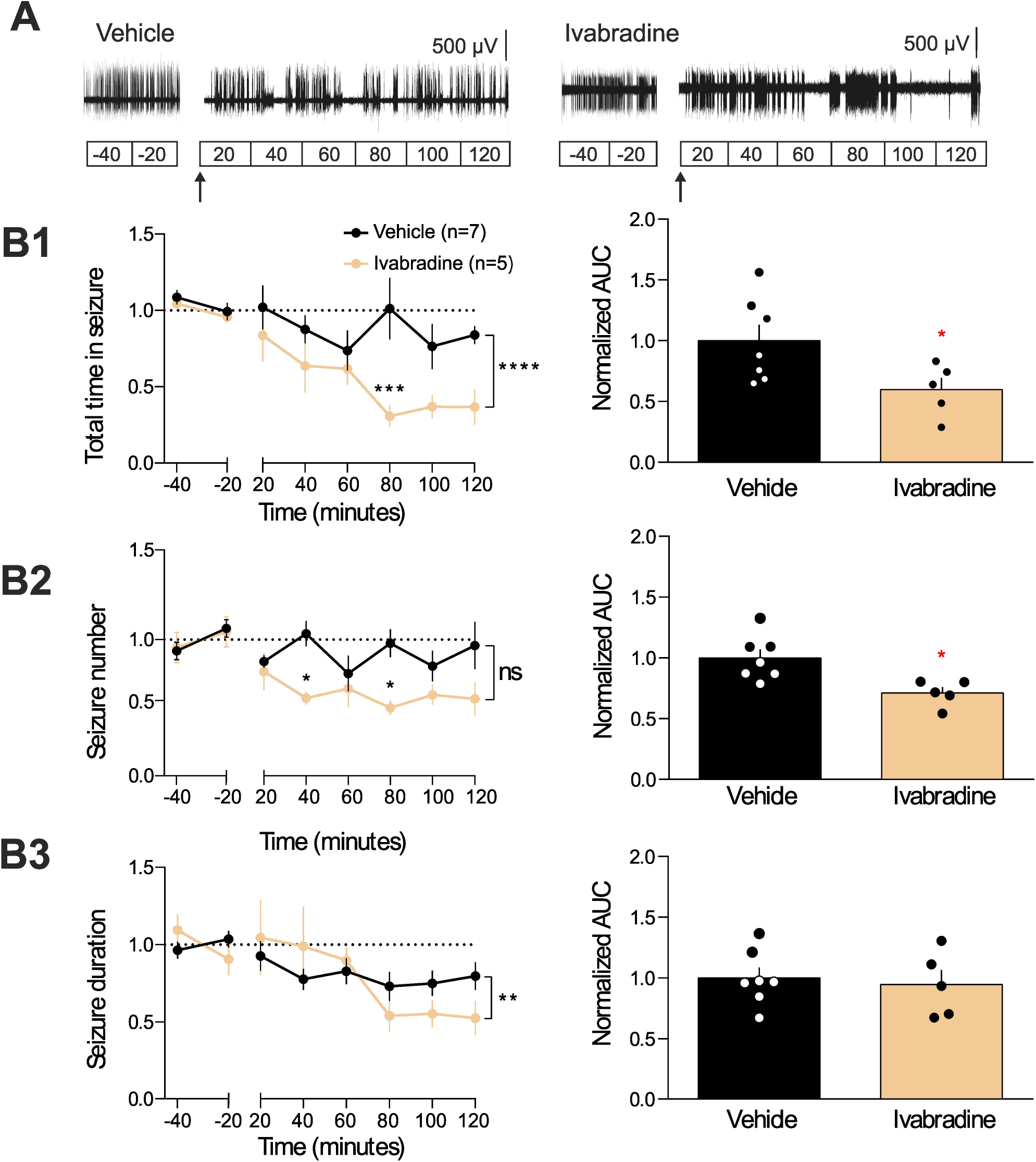
Effect of intrathalamic injection of ivabradine on ASs. **A**. Representative EEG traces of two freely moving GAERS injected in the VB with aCSF or IVA. In **A** and left graphs of **B1-3**, the break in the traces indicates the period of time when the EEG wires were disconnected to allow the exchange of the aCSF with the IVA solution in the injection cannula. aCSF and IVA (6 nmoles/side) were injected using a flow rate of 0.25 µl/min for 2 min. In **B1-3**, data are normalized to the control period. **B1**. Time-response curves (left graph) and area under the curve (AUC) of the whole treatment period (right plot) of the total time spent in seizures for aCSF (black, N=7) and IVA (ochre, N=5) injected animals (*p<0.05, **p < 0.01, ***p<0.005, ****p<0.001). **B2**. Time-response curves (left graph) and AUC of the whole treatment period (right plot) of seizure number of aCSF and IVA treated group (colour-code and number of animals as in **B1**) (*p<0.05, **p < 0.01, ***p<0.005, ****p<0.001). **B3**. Time-response curves (left graph) and AUC of the whole treatment period (right plot) of seizure duration for aCSF and IVA treated animals (colour-code and number of animals as in **B1**).

Microinjection of IVA (6 nmoles) in the CIN abolished ASs even in the first 20 min period after microinjection, and the effect lasted for the entire 2-hour of the post-treatment recording period (**Figure 4A,C1**). The AUC of the total time spent in seizures after IVA (0.04±0.03) microinjection was 96% (F=6.23, DFn=4, DFd=4, p<0,0001) smaller than that of VEH (**Figure 4C1**). Notably, the CIN injection of IVA elicited small-amplitude waves (with no spikes) at 4-7 Hz (**Fig. 4A,B**): these EEG oscillations were not the electrographic expression of ASs since they were not accompanied by motor arrest and the rats kept moving around the cage during this EEG activity. The number of seizures showed a 91% (F=4.57, DFn=4, DFd=4, p<0,0001) decrease in rats treated with IVA (0.09±0.04% compared to VEH) (**Figure 4C2**). Likewise, the mean duration of the seizures was also decreased by 81% (F=1.79, DFn=4, DFd=4,p=0,0002) after IVA administration (0.19±0.08% compared to VEH) (**Figure 4C3**).

**Figure 4.**
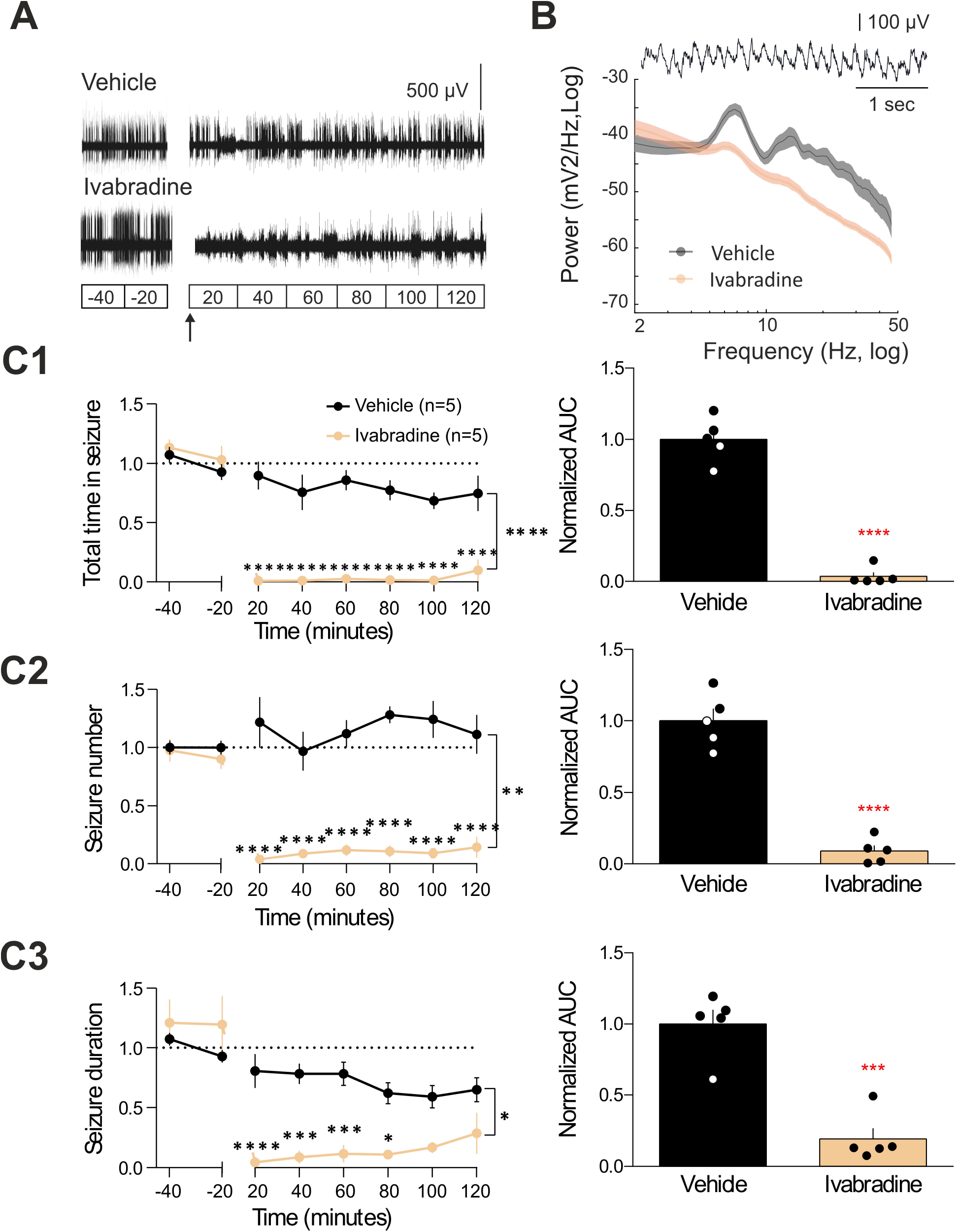
Effect of intra-CIN injection of ivabradine on ASs. **A**. Representative EEG traces of two freely moving GAERS injected in the CIN with aCSF or IVA. In **A** and left graphs of **C1-3**, the break in the traces indicates the period of time when the EEG wires were disconnected to allow the exchange of the aCSF with the IVA solution in the injection cannula. aCSF and IVA (6 nmoles/side) were injected using a flow rate of 0.25 µl/min for 2 min. In **C1-3**, data are normalized to the control period. **B**. Average power spectra of intra-CIN injected vehicle and IVA (number of animals and colour code as in **C1)**. Example of the 4-7 Hz EEG waveform evoked by IVA is shown on top). **C1**. Time-response curves (left graph) and area under the curve (AUC) of the whole treatment period (right plot) of the total time spent in seizures for aCSF (black, N=5) and IVA (ochre, N=5) injected animals (*p<0.05, **p < 0.01, ***p<0.005, ****p<0.001). **C2**. Time-response curves (left graph) and AUC of the whole treatment period (right plot) of seizure number of aCSF and IVA treated group (colour-code and number of animals as in **C1**) (*p<0.05, **p < 0.01, ***p<0.005, ****p<0.001). **C3**. Time-response curves (left graph) and AUC of the whole treatment period (right plot) of seizure duration for aCSF and IVA treated animals (*p<0.05, **p < 0.01, ***p<0.005, ****p<0.001) (colour-code and number of animals as in **C1**).

### Effect of IVA on cortical and thalamic neuron properties

Since the cellular effects of IVA in neurons of key brain areas for AS generation has not been studied either in normal non-epileptic animals and in GAERS, we investigated the ability of this drug to block HCN channel-mediated membrane properties of Wistar cortical layer 5/6 pyramidal neurons (Fig. 5B) and GAERS VB thalamocortical neurons (Fig. 5F) in brain slices. IVA (3 µM) blocked the characteristic depolarizing sag elicited in these neurons by hyperpolarizing current pulses (**Figure 5A,C,E,G**). Furthermore, IVA hyperpolarized the membrane potential of both neuronal types (**Figure 5D,H**) and decreased the number of action potentials evoked by a low-threshold spike in thalamocortical and cortical neurons (VB: control 5.3 ± 0.8, IVA 4.5±0.9, n=10, p<0.05; CIN: control 2.5±0.28, IVA 1.0±0.4, n=6, p<0.05). In contrast, IVA had no effect on the number of action potentials evoked by depolarizing current pulses (VB: control 7.9±0.4, IVA 7.6±0.6, n=10, p>0.05; CIN: control 4.6±0.2, IVA 4.5±0.2, n=6, p>0.05), and the action potential amplitude (VB: control: 71.2±6.8 mV, IVA 71.4±6.5 mV, n=11, p > 0.05; CIN: control: 78.05±4.75, IVA 80.51±5.32, n=6, p>0.05) and threshold (VB: control −51.3±3.3 mV, IVA −51.7±2.7 mV, n=11, p>0.05; CIN: control : −43.7±2.3 mV, IVA −42.6±2.2 mV, n=6, p>0.05), indicating that the effect of this drug is selective on HCN channel-mediated membrane properties in both Wistar CIN cortical layer 5/6 pyramidal and GAERS VB thalamocortical neurons.

**Figure 5.**
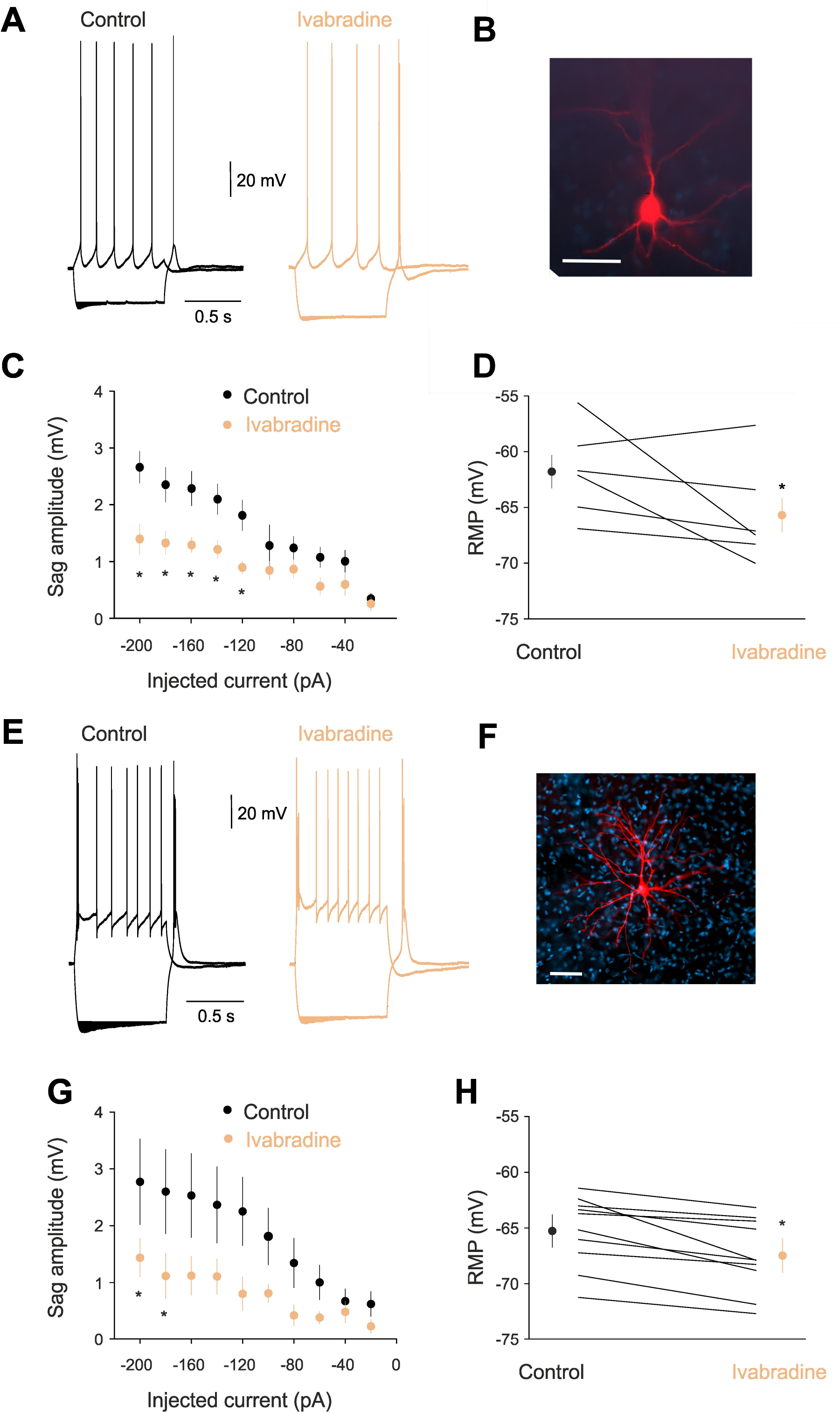
Effect of Ivabradine on the membrane properties of CIN layer 5/6 pyramidal and VB thalamocortical neurons *in vitro*. **A**. Representative voltage responses of a Wistar CIN layer 5 pyramidal neuron to a hyperpolarizing and depolarizing current step (−200 and 80 pA, respectively) before (control) and during application of 30 µM Ivabradine (IVA) (membrane potential: −58 mV). Note the decreased depolarizing sag in the presence of IVA. **B**. Photomicrograph of the cortical layer 5 neuron from which the electrical recordings shown in **A** were made. Scale bar: 50 µm. **C**. Plot of sag amplitude versus injected current show a decrease of the sag in the presence of IVA that is significant for the largest injected currents. **D**. Resting membrane potential in control conditions and in the presence of ivabradine. Large symbols indicate mean ± SEM. **E**. Representative voltage responses of a GAERS VB thalamocortical neuron to a hyperpolarizing and depolarizing current step (−200 and 80 pA, respectively) before (control) and during application of 30 µM Ivabradine (IVA) (membrane potential: −61 mV). Note the decreased depolarizing sag in the presence of IVA. **F**. Photomicrograph of the thalamocortical neuron from which the electrical recordings shown in **E** were made. Scale bar: 50 µm. **G**. Plot of sag amplitude versus injected current show a decrease of the sag in the presence of IVA that is significant for the largest injected currents (p < 0.05). **H**. Resting membrane potential in control conditions and in the presence of ivabradine. Large symbols indicate mean ± SEM.

## Discussion

Our study provides the first demonstration of the potent anti-absence action of systemic administration of the HCN channel blocker, IVA. Moreover, this drug abolishes and reduces ASs when microinjected directly in the CIN and VB, respectively, an action mediated by its ability to decrease I_h_-dependent properties of CIN layer 5/6 pyramidal and VB thalamocortical neurons *in vitro*.

Targeting brain HCN channels *in vivo* has been a major challenge due to the inability of available HCN channel-acting drugs to cross the BBB and provide long-lasting CNS effects. Here, by inhibiting Pgp with ELA,^34,39,40^ we achieved substantial IVA brain concentrations even 2 hours after oral administration to affect physiological brain rhythms, i.e. interictal alpha, theta and gamma waves, and pathological activity, i.e. ASs. Indeed, in the absence of ELA pre-treatment, IVA failed to alter normal and paroxysmal brain oscillations. Notably, the block of Pgp by ELA also allowed for the fast anti-absence action of orally administered IVA observed in this study. In contrast, IVA decreased pharmacologically and electrically induced convulsive seizures without pre-treatment with a drug capable of improving its brain absorption.^31–33^ IVA brain and plasma levels were not measured in these studies, and given our results and previous evidence of IVA inability to substantially penetrate the brain,^27–30^ it is at present difficult to explain the IVA anti-convulsant action reported in the above studies.

The anti-absence effect of IVA injected in the VB and the CIN confirm previous results of the critical role of thalamic and cortical HCN channels in ASs^17–20,25^. The effect of IVA injected in the CIN is markedly stronger than the one following oral administration, while IVA injection in the VB shows a small reduction. In contrast, microdialysis injection in the VB of ZD7288, another HCN channel blocker, has a strong anti-absence action.^25^ Though differences in drug-potency may explain this difference, a much larger portion of the VB is undoutedely affected by continuous 2 hours ZD7288 microdialysis^41^ compared to the more localized 2 min IVA microinjected bolus used in the present study.

Our investigation also provides the first *in vivo* evidence that whole-brain block of HCN channels affects normal brain oscillations, specifically the interictal increase in theta and gamma band power and a shift of the peak of the theta frequency band. Theta and gamma oscillations have been previously linked to I_h_ and ASs^42,43^ and broad changes in cortico-thalamo-cortical firing dynamics induced by blocking I_h_ might underlie both the AS block and changes in interictal oscillations. Moreover, IVA injected in the CIN elicits small-amplitude waves (with no-spikes) at theta frequency (4-7 Hz). The mechanism and patho-physiological significance of all waves induced by systemic and intra-CIN injection of IVA remain to be established.

IVA modulates I_h_-dependent membrane properties of Wistar cortical layer 5/6 pyramidal neurons in the CIN and GAERS thalamocortical neurons in the VB, i.e. the depolarizing sag and the resting membrane potential, leaving other membrane properties unaffected, i.e. action potential threshold and amplitude. As tonic firing was not affected by IVA, the decrease of action potential number evoked by low-threshold spikes of thalamocortical neurons can be explained by the smaller I_h_-tail current in the presence of IVA providing a smaller depolarizing contribution to low-threshold spikes which in turn generate fewer action potentials.

Our study represents the first proof of principle that whole-brain pharmacological block of HCN channels has an anti-absence action, and demonstrate for the first time that reduction of HCN function selectively in the CIN abolishes ASs, i.e. HCN channels of the CIN are necessary for AS generation. HCN1 channels are more abundantly expressed in cortex than thalamus, while HCN2 and HCN4 predominate in thalamocortical neurons.^24,47–51^ Thus, though IVA is a non-selective inhibitor of HCN channel isoforms, it is likely that HCN1 may be the isoform underlying its action in the CIN, whereas its VB effects may involve an interplay between HCN2 and HCN4. Notably, in normal non-epileptic animals, VB-selective knockdown of HCN4 does not elicit ASs, whereas VB-selective HCN2 knockdown, as well as global HCN2^16^ or HCN1^15^ knockout, lead to ASs, suggesting an anti-absence role of both isoforms. In contrast, our previous study^25^ and the present investigation demonstrate that a genetic and pharmacological block of HCN channels in the VB of different mouse and rat models decreases ASs. Possible explanations of these contradictory results include compensatory changes in the full HCN1 and HCN2 KO mice, different species used, different potency of HCN blockers against HCN channel subtypes and diverse efficacy/selectivity of genetic versus pharmacological means of manipulating HCN channels. The alternative interpretation we suggest here is that 1) in normal animals thalamic HCN2 have an anti-absence effect^23^ and thalamic HCN4 a pro-absence action^24,45^ and 2) in epileptic animals there is an increased contribution of HCN4, with respect to HCN2, to the total I_h_ of thalamocortical neurons. Under this hypothesis, in non-epileptic animals VB-selective knockdown of HCN2 elicit ASs and VB-selective knockdown of HCN4 does not.^23,24^ In genetic AS models, a pharmacological or genetic block of all subtypes of thalamic HCN channels would lead to an anti-absence effect.^25^ In support of our hypothesis, i) in thalamic slices of HCN4 KO mice there is a reduction in electrically evoked intrathalamic oscillations (which are considered a proxy of thalamic rhythmic paroxysmal activity)^24^, and ii) an increased I_h_ is present in VB thalamocortical neurons of GAERS^20^ and of normal mice which develop atypical AS following a cortical infarct.^46^ This increased I_h_ may results from enhanced HCN channel expression,^17^ or from changes in the modulation of this current by intracellular messengers (e.g. cAMP)^49^ and neurotransmitters (e.g. noradrenaline).^50^ Notably, our hypothesis makes the testable predictions that in AS models a selective block of thalamic HCN4 and HCN2 channels should have an anti-absence and no effect on ASs, respectively.

In conclusion, acute systemic administration and cortical injection of the HCN channel blocker IVA abolishes genetically determined ASs in a well-established rat model. Selective blockers of HCN channel isoforms, that potentially do not elicit the other changes in EEG oscillations observed here with IVA may represent lead-compounds for future anti-absence drugs.

*‘We confirm that we have read the Journal’s position on issues involved in ethical publication and affirm that this report is consistent with those guidelines*.*’*

### Key Points Box

- Systemic administration of Ivabradine prevents absence seizures by blocking neuronal HCN channels
- Ivabradine injected in the cortical initiation network abolishes absence seizures whereas its anti-absence effect is smaller when injected in the ventrobasal thalamus
- HCN channel blockade by Ivabradine affects membrane properties of cortical layer 5/6 pyramidal and thalamocortical neurons

## Acknowledgements

We wish to thank Dr Nihan Çarçak (Istanbul University) for the help with intracerebral injections. Y. Iacone, T. P. Morais and F. Delicata were supported by a Marie Curie ITN PhD studentship (grant H2020-MSCA-ITN-2016-722053). J. Sandle was supported by the Szeged Scientists Academy under the sponsorship of the Hungarian Ministry of Innovation and Technology (grant FEIF/433-4/2020-ITM_SZERZ). This work was supported by the Ester Floridia Neuroscience Foundation, the Wellcome Trust (Programme Grant 91882 to V.C.), the Eötvös Loránd Research Network (G.T.), the National Research, Development and Innovation Office of Hungary (GINOP-2.3.2-15-2016-00018, Élvonal KKP-20 KKP 133807 to G.T.), the Ministry of Human Capacities, Hungary (20391-3/2018/FEKUSTRAT to G.T.), the Hungarian Scientific Research Fund (Grants NN125601 and FK123831 to M.L.L.) and the Hungarian Brain Research Program (Grant KTIA_NAP_13-2-2014-0014 to M.L.L.).

